# Burst activity plays no role for the field-to-field variability and rate remapping of grid cells

**DOI:** 10.1101/2020.03.28.013318

**Authors:** Michaela Poth, Andreas V.M. Herz

## Abstract

Grid cells in rodent medial entorhinal cortex are thought to play a key role for spatial navigation. When the animal is freely moving in an open arena the firing fields of each grid cell tend to form a hexagonal lattice spanning the environment. Firing rates vary from field to field and change under contextual modifications, whereas the field locations do not shift under such “rate remapping”. The observed differences in firing rate could reflect overall activity changes or changes in the detailed spike-train statistics. As these two alternatives imply distinct neural coding schemes, we investigated whether temporal firing patterns vary from field to field and whether they change under rate remapping. Focusing on short time scales, we found that the proportion of bursts compared to all discharge events is similar in all firing fields of a given grid cell and does not change under rate remapping. Mean firing rates with and without bursts are proportional for each cell. However, this ratio varies across cells. Additionally, we looked at how rate remapping relates to entorhinal theta-frequency oscillations. Theta-phase coding was preserved despite firing-rate changes from rate remapping but we did not observe differences between the first and second half of the theta cycle, as had been reported for CA1. Our results indicate that both, the heterogeneity between firing fields and rate remapping, are not due to altered firing patterns on short time scales but reflect location-specific changes at the firing-rate level.

## Introduction

As a rodent moves through an open arena, the firing fields of each grid cell in the animal’s medial entorhinal cortex (mEC) form an imaginary hexagonal grid tessellating the explored space (Hafting et al., 2005). Despite the striking spatial regularity of these lattices, firing rates vary from field to field by up to an order of magnitude (Diehl et al., 2017; Dunn et al., 2017; Ismakov et al., 2017). These firing-rate variations are stable across time within a session, between repeated sessions, and even after rescaling the arena. When non-spatial cues are altered, however, the field-specific firing rates change even though the firing fields do not shift (Diehl et al., 2017; Ismakov et al., 2017). This phenomenon, known as “rate remapping”, indicates that grid-cell activity represents spatial relations as well as contextual cues.

Grid cells encode spatial information not only at the level of firing rates but also on shorter time scales: During the traversal of a firing field, the spikes of a grid cell tend to occur at successively earlier phases of the theta-band-filtered local field potential (Hafting et al., 2008, Jeewajee et al., 2014). This phase-precession signal is present at the single-run level (Reifenstein et al., 2012; Reifenstein et al., 2014), underscoring its potential role for behavior. High-frequency bursts with discharge rates of up to 300 Hz are often observed (Latuske et al., 2015) and might carry additional information.

Theoretical studies suggest that sensory stimuli can be encoded on fast time scales by modulating the number of spikes that occur within a burst (Kepecs & Lisman, 2003) and such graded burst coding has been revealed in various neural systems (Krahe and Gabbiani, 2004; Eyherabide et al., 2009; Avila-Akerberg et al., 2010). These observations raise the question whether the field-specific firing-rate differences between the multiple firing fields of a given grid cell reflect field-specific spike-train statistics, in particular at short time scales.

To address this question, we reanalyzed grid-cell data from rats foraging in a square enclosure whose wall colors could be changed from black to white (Diehl et al., 2017). We asked whether the burst length or the number of bursts differed when the multiple firing fields of a grid cell were compared. We found that the heterogeneity between firing fields as well as the firing-rate redistribution during rate remapping were unrelated to burst firing. The relative frequency of burst events varied from cell to cell, in agreement with previous findings (Latuske et al., 2015, Csórdas et al., 2019).

Moreover, we asked whether the heterogeneity between firing fields in a grid cell varies throughout the theta cycle. Theoretical and experimental work suggests different computational roles during different theta phases. For example, Sanders et al. (2015) proposed that the first half of a theta cycle is used for the computation of the animal’s present position while future positions are estimated in the cycle’s second half. Our analysis of the theta-phase preference of rate remapping does not support this functional distinction.

Taken together, our results show that burst activity is neither needed to explain the firing-rate variability between the different firing fields of a grid cell nor does it play a role for rate remapping. Both phenomena are fully captured within the classical firing-rate picture. This suggests that the context-dependent modulation of grid-cell activity does not involve inputs that are precisely tuned in time but rather provide a smooth increase or decrease of grid-cell excitability.

## Material and Methods

### Data

We reanalyzed grid-cell data from Diehl et al. (2017), who had recorded 38 grid cells from seven adult male Long Evans rats foraging in a squared enclosure (100 x 100 cm). We excluded two cells from our analysis that showed a very large fraction (more than 33%) of interspike intervals below 4ms. During training and recording, rats explored the arena in blocks of four 10-minute sessions, and were given five minutes between sessions to rest in a box away from the foraging enclosure. The walls of the foraging enclosure were either all black (B) or all white (W) and altered according to a WBB’W’ or BWW’B’ paradigm. The starting color was chosen randomly. Between sessions the enclosure’s floor was cleaned with water. Prior to any electrophysiological recordings, all enclosures were made highly familiar over at least six training days. Behavioral procedures while recording mEC units were identical to training procedures. For details, see the original study.

### Firing-field identification

Similar to Diehl et al. (2017) we constructed firing-rate maps by first summing the total number of spikes that occurred in a given spatial bin (2×2 cm), then dividing by the total amount of time that the rat spent in that bin, and finally smoothing with a Gaussian filter (width: 2.5 cm). To control for possible influences of stationary periods, rate maps were also constructed from data for which the animal’s running speed was above 5 cm/s. Firing-field boundaries were calculated by generating a single reference rate map for the four sessions of one recording block. This map was constructed by averaging the rate maps of the four 10-min sessions. The minimum peak rate required for a field was 2 Hz, the minimum field size was 250 cm^2^.

We then calculated the local maxima of the reference map. Starting from their locations, field boundaries were defined by constructing contours outwards until a threshold value of 0.3 times the individual peak rates was reached. When two fields fused, the threshold value of the higher local maximum was stepwise increased again until the two fields split. The field boundaries derived from the reference map defined the firing field in all four sessions, and all analyses were done for each field in each session. For each such field, the mean field rate was taken as the number of bins in that field divided by the respective dwell time.

### Rate-vector comparisons

To represent the field-specific mean firing rates, we collected their values into one vector for every session. For grid cells with at least 3 firing fields, rate vectors were then compared across sessions using Spearman’s rank correlation. To compare rate vectors to chance, shuffled firing-rate vectors were generated by permuting grid fields such that rate vectors of each grid cell were populated by randomly selected mean firing-field rates.

### Statistical analysis

All analyses were performed in Python 2.7 (RRID: SCR_008394). Specific statistical tests used are stated throughout the text and were taken from Python scipy.stats (RRID: SCR_008058).

### Poisson process

To compare the measured discharge patterns with model spike trains that result in the same rate maps but lack intrinsic bursts, we constructed surrogate spike trains for each recording session of a given grid cell. To do so, we generated rate-modulated Poisson spike trains by using the original rate maps and animal trajectories. As the total number of model spikes in each field might deviate from the measured spike count, we first doubled the firing rates and then randomly drew as many spikes as the original firing field contained. This approach assures that the simulated and the original firing fields have the same mean firing rate.

### Spike-train characteristics

Spike-train autocorrelations and inter-spike interval (ISI) distributions were calculated from binned data. The bin width was 2 ms. For the field-wise comparison of autocorrelations and short ISIs, the spike times of single runs through each firing field were concatenated with a 10 sec interval between the last spike of a run and the first spike of the next run through the same field. A burst was defined as at least two spikes separated by an ISI of less than 10 ms. To compare ISI distributions on time scales relevant for burst firing, the distributions were first normalized in the window from 0 ms to 20 ms.

### Extraction of theta-band oscillations and theta phases from the local field potential

Diehl et al. 2017 recorded not only single-unit activity but also the continuous local field potential (LFP). To extract theta-band oscillations, we filtered the LFP signal with a butterworth bandpass filter (6-11 Hz). The theta phase of a spike was then calculated using the Hilbert transform (taken from Python scipy.signal (RRID:SCR_008058)) of the filtered LFP. We used the convention that 0° denotes the LFP peak. Spike phase histograms were built with 36 bins, each 10° wide, and were normalized by the total number of spikes occurring during the recording.

### Statistical analysis

All analyses were performed in Python 2.7 (RRID: SCR_008394). Specific statistical tests used are stated throughout the text and were taken from Python scipy.stats (RRID: SCR_008058).

## Results

Modifications in contextual cues cause firing-rate changes in rodent grid cells that differ from firing field to firing field and are known as “rate remapping” (Diehl et al., 2017, Ismakov et al., 2017). These changes might reflect changes in the higher-order spike-train statistics or occur independently of fine temporal details in neural activity. To distinguish between these functionally distinct alternatives, we re-analyzed grid-cell data recorded by Diehl et al. (2017). For details, see Material and Methods. These recordings were obtained from the dorsal medial entorhinal cortex (mEC) while rats randomly foraged in a square enclosure (100 x 100 cm) whose walls were black for two sessions (B/B’) and white for two sessions (W/W’). The order of the wall colorations was either BWW’B’ or WBB’W’ and the starting color was chosen randomly. Each block consisted of four 10-minute recording sessions with 5-minute pauses in between, as shown in Figure 1A for a BWW’B’ session. The location of grid fields (see also Material and Methods: Firing-field identification) remained constant but the field-specific firing rates changed (Figure 1B). This can readily be seen by calculating the mean firing rate for each firing field and entering these values into a rate vector (see Material and Methods). Comparing these vectors across the four sessions of each cell (Figure 1C) shows that the firing rates of corresponding fields are similar between sessions with matching colors but not across sessions with non-matching colors, as reported by Diehl et al. (2017).

**Figure 1:**
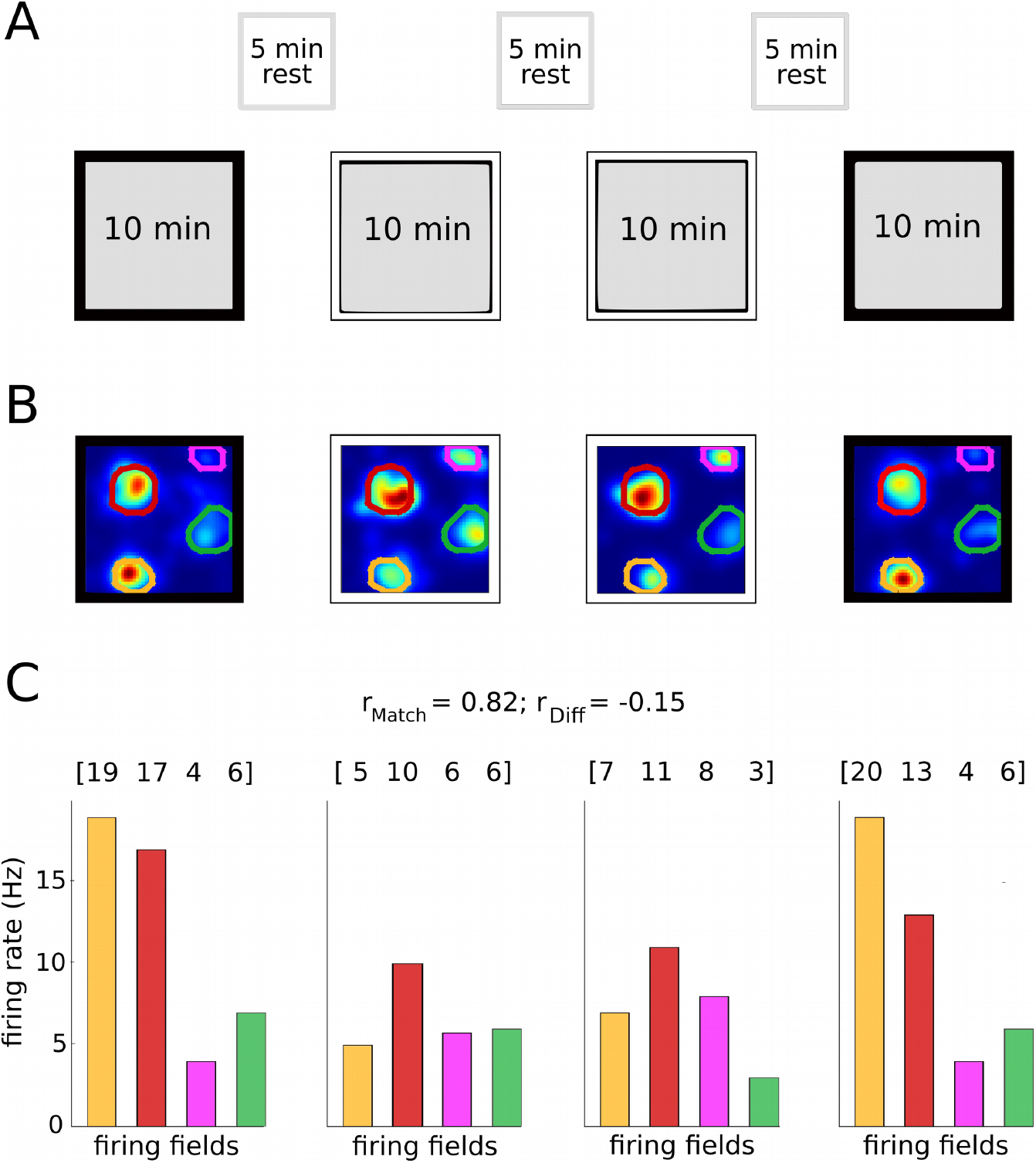
Remapping of grid-cell firing rates. (A) Schematic drawing of the experimental paradigm. With interleaved resting periods, the animal explored a square box whose wall colors were altered between black and white according to a BWW’B’ or WBB’W’ sequence. (B) Firing-rate maps of an example grid cell that was recorded across all four conditions. The individual firing fields are encircled by a colored line. Shape and position of each firing field is almost identical in all four conditions. (C) Mean firing rates, in matching colors, for the four fields shown in (B). The Rate Vectors (RVs) above the color bars represent the mean firing rates within each grid field. The average Spearman correlation of the RVs is high between matching conditions but low across different conditions.

### Heterogeneity between firing fields is not the result of altered burst firing

Consistent with previous reports (Diehl et al., 2017, Dunn et al., 2017, Ismakov et al., 2017), we observed that within a given environment grid-cell firing rates varied strongly from field to field. To quantify this finding, we calculated the coefficient of variation (CV) of the mean in-field firing rates from the firing-rate maps of each cell and session. Across these samples, we obtained an average CV of 0.40 (Figure 2A).

**Figure 2:**
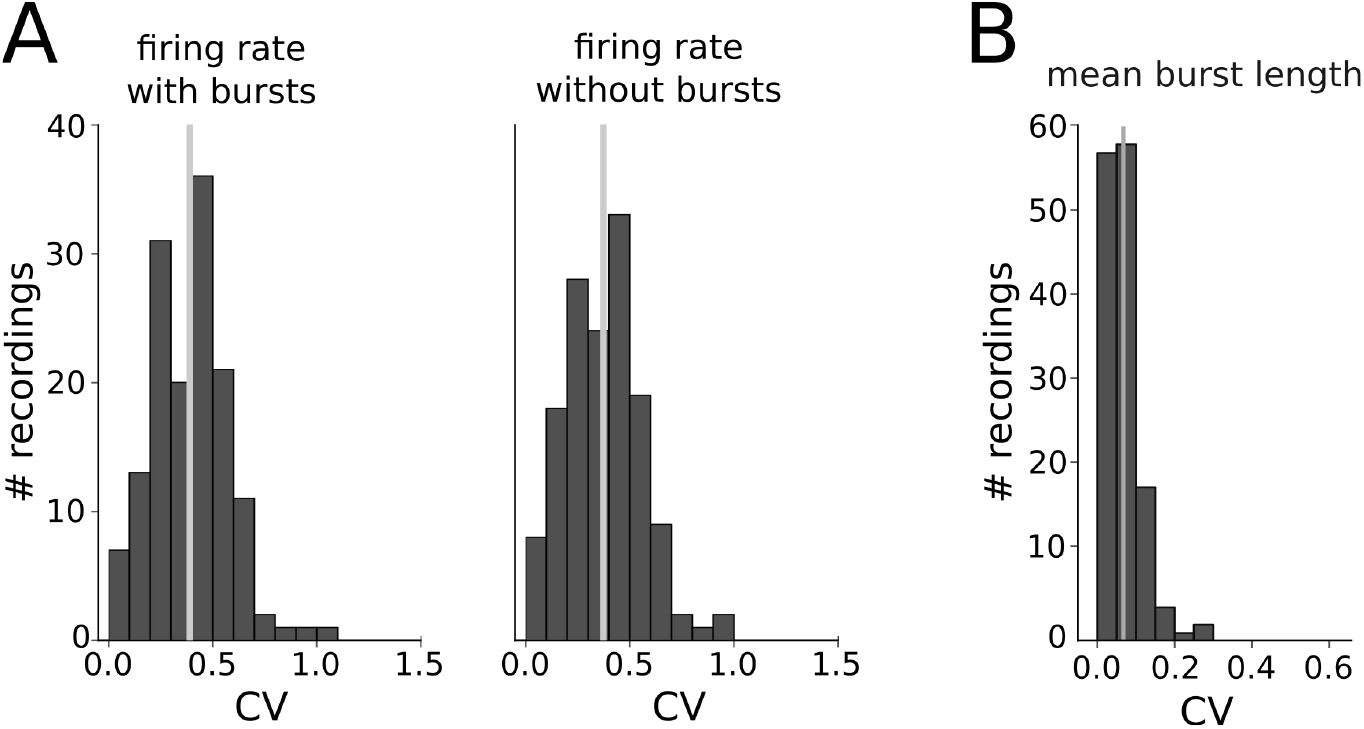
Heterogeneity of firing fields is not caused by burst activity. (A) Population results for the firing-rate variability. For each grid cell and recordings session, firing fields were identified and mean discharge rates were computed field-by-field. Their variability was measured in terms of the coefficient of variation (CV) and averaged across sessions and cells. The left panel is based on the original spike data. For the second panel burst-like discharges were discarded by removing all spikes with a preceding inter-spike interval (ISI) of less than 10 ms. (B) Population results for the burst-length variability, calculated as the CV of the number of spikes within a burst.

If this variability resulted from differences in burst firing, the CV should decrease strongly after bursts have been removed from the analysis. In a next step, we therefore excluded all spikes following another spike with an inter-spike interval (ISI) of less than 10ms (“without burst”) so that only isolated spikes and the first spike of each burst remained for the analysis. The CV distributions of the mean in-field firing rates without bursts had an average CV of 0.39 and were indistingishable from those with bursts (two-sample t-test: statistic=0.71, p-value=0.48). Repeating the same analyses with a cut-off of 6ms or 8ms did not change the results (data not shown). Furthermore, for each recording the mean burst length was highly similar across firing fields (average CV: 0.06) (Figure 2B). These data demonstrate that burst firing does not explain the heterogeneity in firing between the firing fields of each grid cell and session.

### Rate remapping does not hint at changes in burst firing

Although the firing-field variability within one enclosure does not hint at differences in burst firing, burst firing might change under rate remapping. To tackle this possibility, we performed two analyses to detect similarities or differences in burst behavior. First, we compared the four ISI histograms of a given grid cell and focused on ISI-values relevant for burst firing (ISI<20ms). We observed that ISI histograms were similar across different conditions but differed from cell to cell, as shown by the two example cells in Figure 3A. We quantified this finding by calculating the Kullback–Leibler (KL) divergence between the normalized probability density functions of the ISI distributions for WW (W/W’), BB (B/B’), and BW (B/W, B/W’, B’/W and B’/W’) and compared the distributions using a Kolmogorov-Smirnov (KS) test. This test showed that the ISI distributions were similar in the white and black boxes (WW vs. BB: KS test: statistic=0.18, p-value=0.62; WW vs. BW: KS test: statistic=0.14, p value=0.63; BB vs. BW: KS test: statistic=0.15, p-value=0.51). In contrast, the ISI histograms differed from cell to cell. This can be seen by comparing the KL divergence across environments with the distribution across cells (KS test: statistic=0.45, p-value=8.98e-11). We then wondered whether the same holds true for spike-train autocorrelations. In a first step, we calculated the Pearson-Correlation-Coefficient of matching (median correlation coefficient for WW across all neurons: r=0.85; for BB: r=0.85) and non-matching colors (r=0.83). A KS test showed that the three distributions were similar (WW vs. BB: statistic=0.18, pvalue=0.62; WW vs. BW: statistic=0.18, pvalue=0.29; BB vs. BW: statistic=0.15, pvalue=0.50), as depicted in Figure 3B for same two example cells as in Figure 3A. As a control, we created sets of surrogate data. To this end, we combined spike-train autocorrelations from two different cells and calculated the Pearson-Correlation-Coefficient of matching (median correlation coefficient for WW surrogate: r=0.63; for BB surrogate: r=0.69) and non-matching colors (BW surrogate r=0.64). The KS test showed a significant difference of the WW/WW surrogate (statistic = 0.53, p-value: 7.29e-05), BB/ BB surrogate (statistic = 0.50, p-value = 2.20e-04), and BW/BW surrogate (statistic = 0.53, p-value = 2.07e-07). We conclude that there are significant cell-to-cell differences in the ISI distributions and spike-train autocorrelations but no contextual changes when all firing fields of a grid cell are considered together.

**Figure 3:**
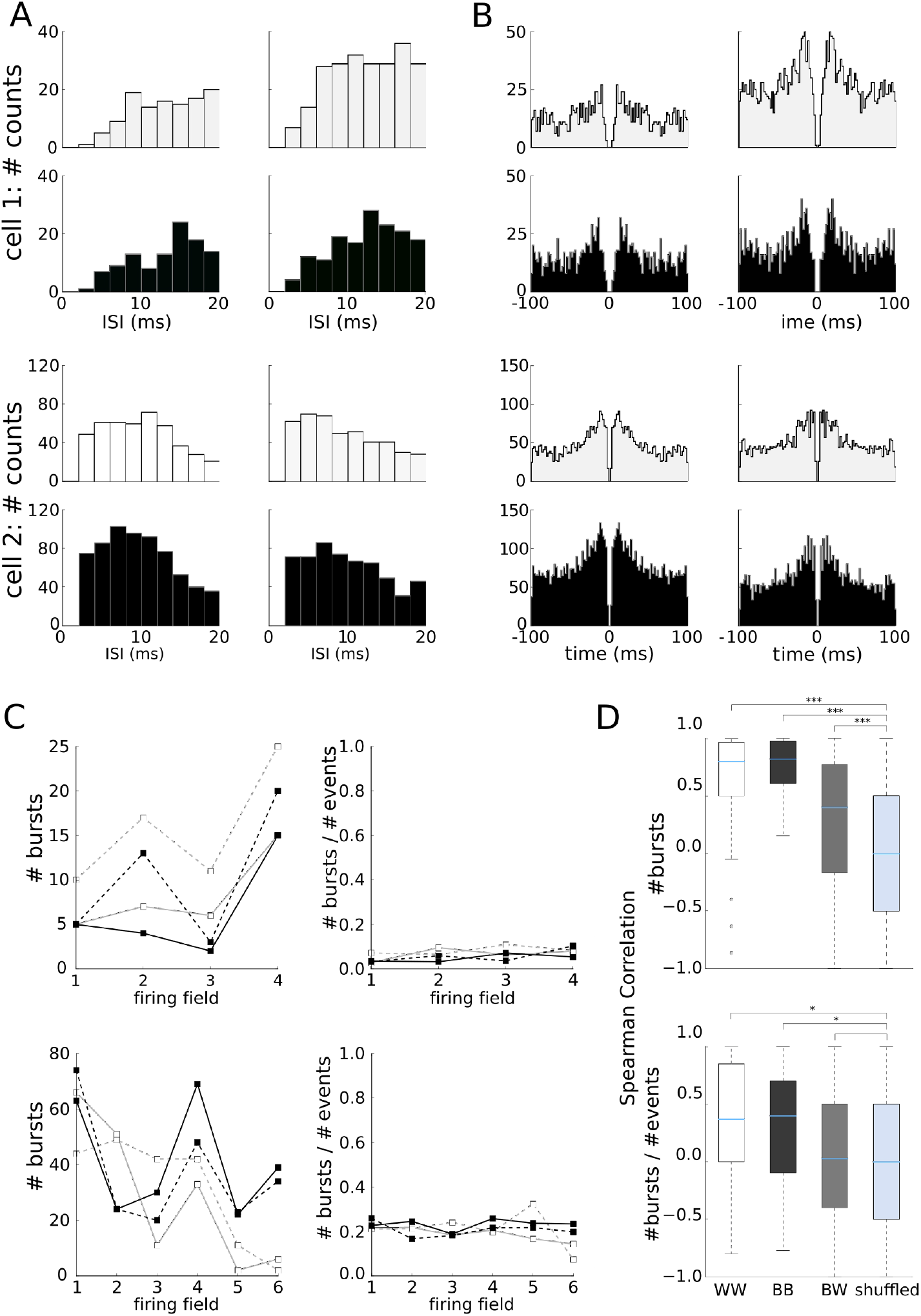
Rate remapping does not hint at changes in burst firing. (A) ISI histograms and (B) autocorrelations of two example cells. The white and black colors of the filled areas mark the colors of the respective enclosure walls. (C) Number of bursts per firing field (left) and relative number of bursts (right). Black/white enclosures are indicated by lack/grey lines, respectively. Line style denotes the first (solid) and second (dashed) session in that enclosure. (D) Population results of the absolute and relative burst numbers across all cells, as quantified by Spearman Correlations for the specific wall combinations WW, BB and BW, as well as for shuffled labels. Blue lines indicate the population median.

Next, we evaluated whether the redistribution of mean in-field firing rates across grid fields results from differences in bursting behavior. To this end, we counted the absolute and relative number of bursts within each grid field and entered these values into a vector for each grid cell and each 10-min session (Figure 3C). By comparing the vectors across sessions, we confirmed that the absolute number of bursts were similar between sessions with matching box colors (median Spearman’s rank correlation for WW: 0.80; for BB: and 0.82) as shown in Figure 3D. For non-matching box colors we observed a weaker correlation (median Spearman’s rank correlation: 0.40). Label shuffling led to a median Spearman’s rank correlation of 0.00. This demonstrates that the correlations between sessions in matching and non-matching box colors are significant (KS test for WW: statistic=0.54, p-value=1.16e-09; for BB: statistic=0.67, p-value=1.03e-14; for BW: statistic=0.21, p-value=4.14e-06). In contrast, the relative number of spikes shows a low median Spearman’s rank correlation for the matching and non-matching cases (median Spearman’s rank correlation for WW: 0.37; for BB: 0.39; for BW: 0.03) as depicted in Figure 3D. Label shuffling indicates that the correlations are weakly significant (KS test for WW: statistic: 0.25, p-value: 0.02; for BB: statistic: 0.24, p-value: 0.02) for matching colors and not significant for non-matching colors (KS test: statistic=0.11, p-value=0.08). The relative number of spikes rather fluctuates around one specific value, which varies from cell to cell. We also did not see effects that depend on the order of box-color changes. This indicates that the redistribution of the firing rates in the color-change paradigm is not the result of altered burst firing.

### Mean in-field firing rates with and without bursts are proportional for each cell

As changed burst firing is neither required to explain rate remapping nor the firing-rate variability across fields, we expected that in-field firing rates with and without bursts were correlated. Indeed, we observed a linear relation between the firing rate with and without bursts (Figure 4A). For each cell, a linear regression line fits the data well (p-values < 1.35e-05). The slopes vary between 0.62 and 0.95 and the intercepts between −0.20 and 0.57 (Figure 4B). Since the distributions are similar for black and white environments (Wilcoxon-rank sum test: slope: statistic=293.0, p-value=0.53, intercept: statistic=236.0, p-value=0.43), we pooled the data for each cell across environments. However, the variability in the slopes did not appear compatible with a simple Poisson process. To quantify this finding, we simulated spike trains generated with inhomogeneous Poisson processes based on the individual firing-rate maps and animal trajectories (Materials and Methods) and repeated the previous analysis. To create model spike trains without “bursts”, we removed all spikes following an inter-spike interval of less than 10ms. We again found a linear relation between the mean in-field firing rate with and without “bursts” (Figure 4C). The distribution of the slopes ranged from 0.75 to 0.94 (Figure 4D) and differed from the original data (Wilcoxon-rank sum test: slope: statistic=6.0, p-value=2.79e-07). This was to be expected as for a Poisson process, the relation between the number of “bursts” and the number of all events is similar in each firing field.

**Figure 4:**
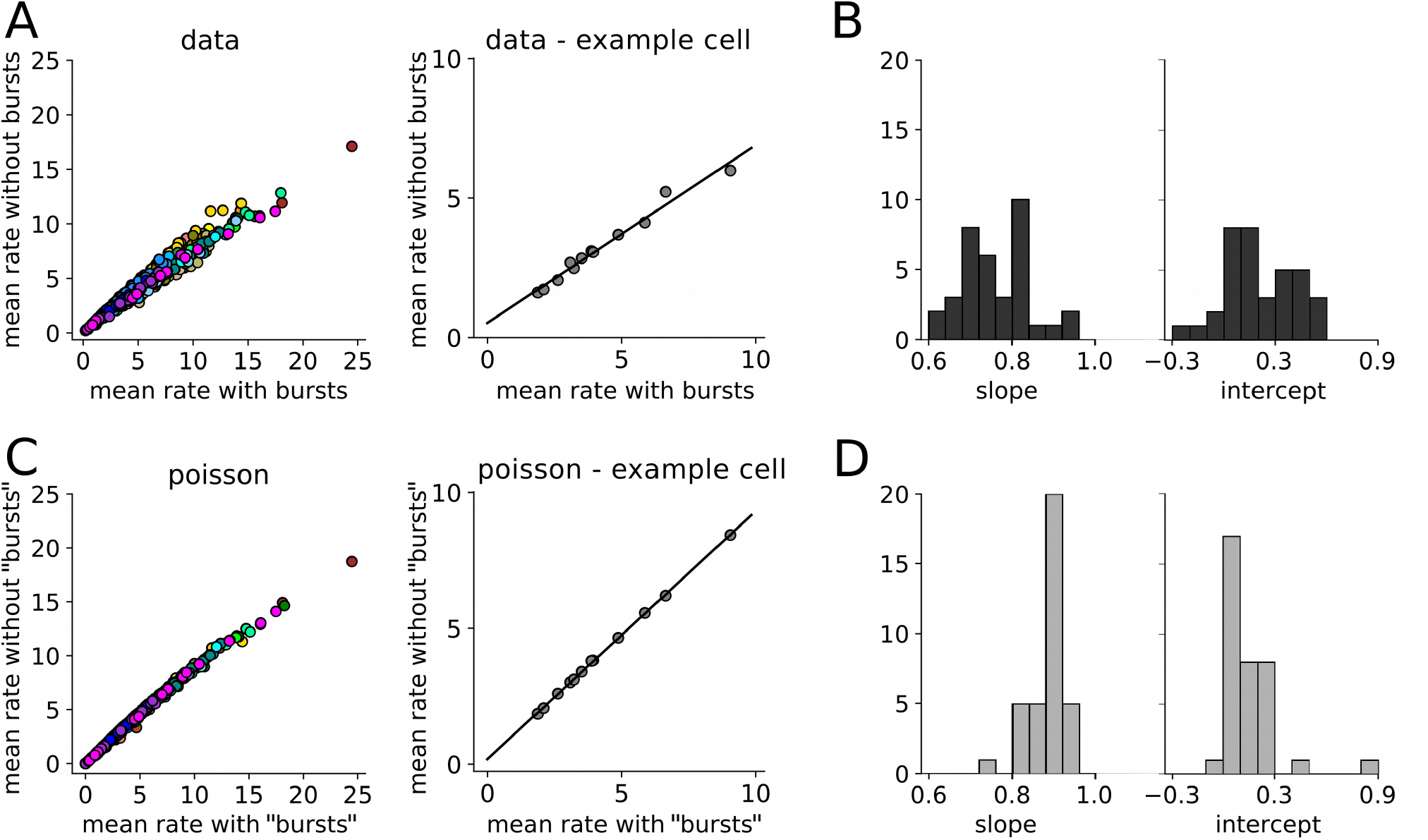
Mean firing rates with and without bursts are proportional. (A) Each dot represents the mean firing rate in a firing field calculated with all spikes (“with bursts”) on the x-axis and after spikes with a previous ISI < 10 ms were removed (“without bursts”) on the y-axis. Left panel: Population results. Firing fields of a specific cell have the same color. Right panel: One example cell, together with the linear regression line. (B) Population results from measured grid cells: Distribution of the linear regression slopes (left column) and distribution of the linear regression intercepts (right column). (C) Model firing rate maps were simulated with inhomogeneous Poisson processes that were based on the original firing rate maps and movement patterns. The firing field sizes and positions as well as the number of spikes per firing field are exactly the same than in the original data. Left panel: Population results. As in (A), the firing fields of a specific cell have the same color. Right panel: One example cell, together with the linear regression line. (D) Population results as in (B) but now from the Poisson model.

These findings underscore that grid-cell firing deviates from Poisson spiking but do not hint at a particular role of burst events.

### Rate remapping is not reflected in the local field potential

In line with previous work (Hasselmo et al., 2002; Sanders et al., 2015), Sanders et al. (2018) provided evidence that under hippocampal rate remapping, the two halves of the theta cycle have different functions for place cells in CA1, but not in CA3. For each cell, these authors compared place fields in which rate remapping was observed. On a place-field-by-place-field basis, they defined a “low-rate condition” as the condition for which the cell’s firing rate (in that particular field) was lower than in the other condition, named “high-rate condition” (for that field). In the low-rate condition, place cells in CA1 tended to spike during the second half of the theta cycle while the first half was preferred in the high-rate condition. This type of theta-phase dependence was not observed in CA3. The medial entorhinal cortex, populated by grid cells, projects both directly and indirectly to CA1 and CA3. This raises the question whether grid-cell firing under remapping is reminiscent of place-cell firing in CA1 or place-cell firing in CA3 or shows yet another behavior.

Before we address this question, let us first ask whether for a given environment, spike phases differ between grid fields with higher rate and fields with lower rate. To this end, we constructed – for each grid cell and recording – a polar histogram of the spikes occurring in the grid field with the highest mean firing rate and of the spikes occurring in the grid field with the lowest mean firing rate. We then calculated the (circular) difference of the circular means of both histograms for each recording session (Figure 5A). Across cells and sessions, circular statistics gives a mean difference of −7° with a standard deviation of 52° (Figure 5B, left panel). These data suggest that field-to-field differences in the firing rates of a given grid cell are not mirrored in phase differences relative to the LFP.

**Figure 5:**
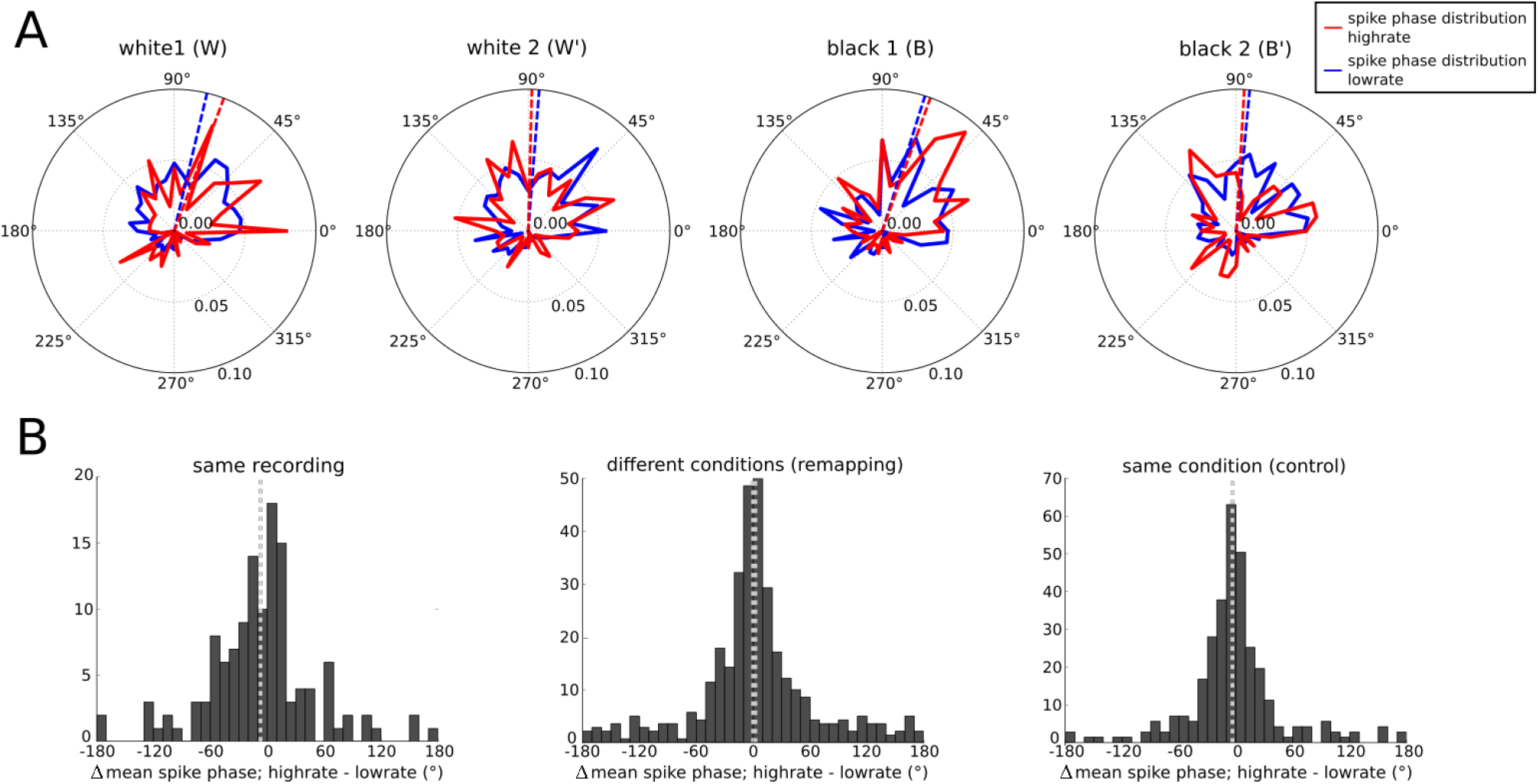
Spike phases do not differ between high and low firing rates. (A) For one example cell, distributions of spike phases relative to the theta-band oscillation in the local field potential (LFP) are shown for the four environments (W, W’, B, B’). Red (blue) curves represent the spike phases in the firing field with the highest (lowest) mean firing rate. Dashed lines mark the circular means of the two distributions. (B) The differences between the circular mean of the spike phase distribution in the high rate condition and the low rate condition are shown. For the left panel the high/low rate condition is defined as in Figure 5A, i.e., as the firing field with the highest/lowest mean firing rate within each recording. All four environments are treated independently and the differences between the two circular means (high rate minus low rate) are collected across cells and environments. In the middle panel we compared spike phases from the same firing fields under rate remapping, as Sanders et al. (2018) did for hippocampal place cells. Accordingly, high/low-rate condition is defined as the condition with the higher/lower mean firing rate for that particular firing field. All scenarios (W/B, W/B’, W’/B, and W’/B’) are considered across all cells. In the right panel we compare, as a control, the recordings from the two white/black environments of each cells. The two possible combinations (W/W’ and B/B’) are pooled across all cells. In the third panel we considered the recording from the white and the black environments of each cell. As the position of each firing fields is the same in all four recording sessions, we compared the spike phase distributions for each firing field in W/B, W/B’, W’/B, and W’/B’. We defined the high/low-rate condition as the firing field with the higher/lower mean firing rate. For each firing field the circular means of the phase distributions were calculated. The distribution of the differences of the circular means are shown across cells. In all three panels the light grey dotted line denotes the circular mean of the respective distribution.

Still, rate remapping could cause phase shifts. Following Sanders et al. (2018), we compared high-rate and low-rate conditions across differently colored environments (BW). Circular statistics did not reveal significant differences in the theta-phase preferences between the high-rate and the low-rate conditions (mean = 1°, std = 55°), as shown in the middle panel of Figure 5B. As a control, we also compared high-rate vs. low-rate conditions from the first and second recording with the same coloring (WW or BB). Here, we obtained a mean of −4° (std = 43°), see the right panel of Figure 5B. We conclude that mEC grid cells do not show the firing-rate-dependent phase-preferences of CA1 place cells but rather the characteristics of CA3 cells. This suggests that the behavior exhibited by CA1 place cells is not caused by mEC grid-cell inputs.

## Discussion

Grid cells encode spatial information not only in their firing-rate based activity fields but also at a finer temporal scale via theta-range phase precession (Hafting et al., 2008). In addition, grid cells represent contextual information through field-to-field variations in their firing rates (Diehl et al., 2017; Ismakov et al., 2017). Whether these variations are due to differences in the higher-order spike statistics, such as burst firing, or simply result from different activity levels has not been addressed in the literature. To tackle this question, which is key for a comprehensive understanding of the grid-cell code, we investigated the fine-scale temporal behavior of grid cells field-by-field and studied its potential impact on the firing-rate variability.

It has been suggested that burst duration might encode information (Kepecs and Lisman., 2003), and indeed, burst duration coding is present in other neural systems (Eyherabide et al., 2009, Avila-Akerberg et al., 2010). However, in the grid-cell data analyzed in the present study, the number of spikes per burst does not vary significantly between firing fields. Furthermore, we found that the appearance of bursts does not influence the heterogeneity between firing fields. We also did not find differences in the ISI distributions (ISI < 20 ms) and the autocorrelations of each grid cell within the same or across differently colored enclosures. Furthermore, although the absolute number of bursts varied from field to field, the ratio between bursts and all events was constant between fields and recording sessions. This indicates that individual grid cells have a specific bursting behavior and the redistribution of the firing rates across their grid fields is not the result of modified burst firing. This finding is supported by the observation that in all firing fields of a given grid cell the mean firing rates with and without bursts are proportional. Different cells, however, differ in their burst behavior. This is consistent with the existence of bursty and non-bursty grid cells (Latuske et al., 2015, Csórdas et al., 2019). It remains an open question whether the burstiness of a grid cell differs in novel versus familiar environments.

Our analysis shows that neither the heterogeneity of the firing fields of a given grid cell nor the redistribution of its firing rates under contextual changes is the result of altered burst firing. These results are in line with findings in CA1 and CA3, which demonstrate that hippocampal rate remapping is not the result of modulations in burst firing (Sanders et al., 2018). We have also shown that neither the heterogeneity of the firing rates nor rate remapping leads to changes in the preferred spike phases. This is in line with results shown for rate remapping of place cells in CA3 but not with the results of place cells in CA1 (Sanders et al., 2018). The grid cells analyzed by Diehl et al. (2017) reside in superficial mEC layers which directly project to CA1 and CA3. In contrast to phase precession phenomena (Schlesiger et al., 2015), the preferred spike phases during rate remapping in

CA1 can thus not be inherited form grid cells. More generally, our findings suggest that the field-to-field variability of grid-cell firing and its context-dependent modulation are not caused by precisely timed inputs but rather by a gradual increase or decrease of grid-cell excitability.

## Acknowledgements

This work was supported by the Deutsche Forschungsgemeinschaft through the Research Training Group 2175 (Perception in Context and its Neural Basis). We thank G.W. Diehl, S. Leutgeb, and J.K. Leutgeb for making data from Diehl et al. (2017) available; and S. Häusler and C. Leibold for stimulating discussions.

## References

Ávila-Åkerberg O, Krahe R, Chacron MJ (2010) Neural heterogeneities and stimulus properties affect burst coding in vivo. Neuroscience 168:300–313.

Csordás DE, Fischer C, Nagele J, Stemmler M, Herz AVM (2019) Spike afterpotentials shape the in-vivo burst activity of principal cells in medial entorhinal cortex. bioRxiv:10.1101/841346.

Diehl GW, Hon OJ, Leutgeb S, Leutgeb JK (2017) Grid and nongrid cells in medial entorhinal cortex represent spatial location and environmental features with complementary coding schemes. Neuron 94:83–92.

Dunn B, Wennberg D, Huang Z, Roudi Y (2017) Grid cells show field-to-field variability and this explains the aperiodic response of inhibitory interneurons. arXiv:1701.04893v1

Eyherabide HG, Rokem A, Herz AVM, Samengo I (2009) Bursts generate a non-reducible spike-pattern code. Frontiers in Neuroscience 3:8–14.

Hafting T, Fyhn M, Molden S, Moser M-B, Moser EI (2005) Microstructure of a spatial map in the entorhinal cortex. Nature 436:801–806.

Hafting T, Fyhn M, Bonnevie T, Moser MB, Moser EI (2008) Hippocampus independent phase precession in entorhinal grid cells. Nature 453:1248–1252.

Hasselmo ME, Bodelón C, Wyble BP (2002) A proposed function for hippocampal theta rhythm: separate phases of encoding and retrieval enhance reversal of prior learning. Neural Computation 14:793–817.

Ismakov R, Barak O, Jeffery K, Derdikman D (2017) Grid cells encode local positional information. Current Biology 27:2337–2343.

Jeewajee A, Barry C, Douchamps V, Manson D, Lever C, Burgess N (2014) Theta phase precession of grid and place cell firing in open environments. Phil. Trans. R. Soc. B 369:20120532

Kepecs A, Lisman J (2003) Information encoding and computation with spikes and bursts. Network: Computation in Neural Systems 14:103–118.

Krahe R, Gabbiani F (2004) Burst firing in sensory systems. Nature Reviews Neuroscience 5:13–22.

Latuske P, Toader O, Allen K (2015) Interspike intervals reveal functionally distinct cell populations in the medial entorhinal cortex. Journal of Neuroscience 35:10963–10976.

Reifenstein ET, Kempter R, Schreiber S, Stemmler MB, Herz AVM (2012) Grid cells in rat entorhinal cortex encode physical space with independent firing fields and phase precession at the single-trial level. Proc. Natl. Acad. Sci. USA 109:6301–6306.

Reifenstein E, Stemmler M, Herz AVM, Kempter R, Schreiber S (2014) Movement dependence and layer specificity of entorhinal phase precession in two-dimensional environments. PLoS one 9:e100638

Sanders H, Rennó-Costa C, Idiart M, Lisman J (2015) Grid cells and place cells: an integrated view of their navigational and memory function. Trends in Neurosciences 38: 763–775.

Sanders H, Daoyun J, Sasaki T, Leutgeb JK, Wilson MA, Lisman JE (2018) Temporal coding and rate remapping: Representation of nonspatial information in the hippocampus. Hippocampus 2018:1–17.

Schlesiger MI, Cannova CC, Boublil BL, Hales JB, Mankin EA, Brandon MP, Leutgeb JK, Leibold C, Leutgeb S (2015) The medial entorhinal cortex is necessary for temporal organization of hippocampal neuronal activity. Nature Neuroscience 18:1123.

